# Estradiol promotes habituation learning via an unidentified target, bypassing the suppressive effects of established Estrogen Receptors

**DOI:** 10.1101/2025.02.03.636196

**Authors:** Andrew Hsiao, Isabelle Darvaux-Hubert, Dominique Hicks, Emilie Joux, Sarah De Freitas, Emeline Dracos, Jeanne Lizé, Julien Perrichet, Dominique Baas, Owen Randlett

## Abstract

Habituating to the constant stimuli in the environment is a critical learning process conserved across species. We use a larval zebrafish visual response to sudden darkness as a model for studying habituation learning, where zebrafish reduce their responses to repeated stimulations. In this paradigm, treatment with estradiol strongly increases learning rate, resulting in more strongly suppressed responses. We used mutant lines for the Estrogen Receptors (*esr1, esr2a, esr2b, gper1*) in an attempt to identify the receptor(s) mediating these effects. These experiments failed to identify a necessary receptor (or combination of receptors). Surprisingly, *esr1, esr2a*, and *gper1* mutants showed weak but consistent increases in habituation, indicating these receptors suppress habituation learning. These experiments demonstrate that estradiol is a complex modulator of learning in our model, where the learning-promoting effects are mediated by an unidentified estradiol target, and the classical Estrogen Receptors act in competition to subtly suppress learning.

## Introduction

A primary function of the brain is to learn from experiences and adjust behavior in response. One aspect of learned behaviour involves sharpening attention and behavioural resources toward salient cues by ignoring irrelevant background stimuli. For instance, it may be critical to recognize the alarm calls of a nearby animal, whereas continually registering the steady hum from distant traffic is far less important. The capacity to reduce responses to repetitive, non-essential stimuli is known as habituation, which is considered the simplest form of learning and memory (***Rankin et al., 2009***).

We study a paradigm for long-term habituation where larval zebrafish reduce their responsiveness to sudden pulses of whole-field darkness, or dark flashes (DFs) (***Wolman et al., 2011***; ***Randlett et al., 2019***; ***Lamiré et al., 2023***). We recently reported that multiple hormonal signaling pathways show strong modulation of habituation learning performance, including melatonin, progesterone, and estrogen (***Lamiré et al., 2023***). The ability of these signaling pathways to modulate learning is consistent with previous results in other systems and paradigms (***Nilsson and Gustafsson, 2002***; ***Naderi et al., 2020***; ***Dillon et al., 2013***; ***Rawashdeh et al., 2007***; ***Jilg et al., 2019***; ***El-Sherif et al., 2003***; ***Barros et al., 2015***), and may be an important mechanism to shift learning and memory performance or strategies based on biological rhythms or external fluctuations like seasons, weather or the day/night cycle.

In this project we have focused on estrogen signaling. We identified multiple estradiol analogs which strongly increased habituation learning when bath applied at 5-10μM doses (ethinyl estradiol, estradiol valerate, and hexestrol, ***Lamiré et al., 2023***). 17β-estradiol (here referred to as estradiol) is the most potent and biologically active form of estrogen, and is used in a variety of clinical contexts including contraception, hormone replacement therapy, and feminizing hormone therapy (***Kuhl, 2005***; ***Unger, 2016***; ***Farkas et al., 2022***).

Our discovery of a role for estradiol in promoting habituation learning is not surprising, as it has well documented effects on other learning and memory processes (***Frick, 2015***). This has been most extensively characterized in the hippocampus, where estradiol promotes behavioural performance and the cellular/circuit hallmarks of hippocampal plasticity, including Long-term potentiation (LTP) and modulation of dendritic spine density (***Iqbal et al., 2024***; ***Luine, 2014***; ***Finney et al., 2020***; ***Nilsson and Gustafsson, 2002***). While the role of estradiol in habituation is less well explored, it has previously been shown to increase memory retention for olfactory habituation in mice (***Dillon et al., 2013***), indicating it plays conserved roles in plasticity regulation across paradigms.

Estradiol signals via two established classes of Estrogen Receptors (ERs): the ligand-activated transcription factors ERα, and ERβ, and the seven-transmembrane G-protein coupled receptor Gper1. ERα/β are thought to mediate the long-term “genomic” effects of estrogens through transcriptional activation of target genes, and thus are typically termed nuclear ERs. Estradiol also has acute effects on physiology – often called “non-genomic”, or “membrane-initiated”. These acute effects are thought to be mediated largely by the G protein coupled receptor Gper1, which signals via multiple G-proteins, and potentially epidermal growth factor (EGF) receptor transactivation (***Prossnitz and Barton, 2023***; ***Revankar et al., 2005***; ***Filardo et al., 2000***). In this way, Gper1 signalling impacts multiple core second-messenger systems, including: adenylyl cyclase, ERK, PI3K-Akt, and nitric oxide synthase. There is evidence from receptor-specific pharmacology and genetic/mutant experiments in mice for a role of all of these receptors in hippocampal plasticity (***Finney et al., 2020***; ***Koitm*ä*e et al., 2023***; ***Briz et al., 2015***).

Pharmacological experiments in adult zebrafish indicate that nuclear ERs are involved in the consolidation of object recognition memory, while Gper1 is involved in the consolidation of object placement memory (***Naderi et al., 2020***). Therefore, both classes of receptors are good candidates for plasticity regulation in zebrafish. However, we are unaware of any previous studies using mutant lines to test for the function of zebrafish ERs in learning or memory.

In this project we aimed to identify the relevant ER(s) mediating the effects of estradiol on habituation using mutant alleles. Zebrafish have single gene encoding ERα (ER1, *esr1*) and Gper1 (*gper1*), and two homologs of ERβ: ER2a (*esr2a*) and ER2b (*esr2a*) (***Romano et al., 2017***; ***Menuet et al., 2002***). We found that none of these mutants were insensitive to estradiol’s effects, indicating that estradiol acts in this context via an alternative receptor or pathway. Surprisingly, our experiments found that mutants for *esr1, esr2a*, and *gper1* actually habituate more than their sibling controls. While the effect size is small and behavioural-genetic experiments can be variable, these data indicate that these ERs actually act to inhibit habituation learning, rather than mediating the habituation-promoting effects of estradiol that we observe pharmacologically.

## Results

### Estradiol increases visual habituation learning

In response to a sudden global darkening stimulus, which we refer to as a dark flash (DF), larval zebrafish execute an “O-bend” maneuver, characterized by a deep “O”-shaped bend and a high-amplitude turn (***Burgess and Granato, 2007***, ***Figure 1***a). Habituation learning manifests as a progressive reduction in response to repeated stimuli, and this learning can be retained for seconds/minutes, or hours/days for short-, and long-term habituation, respectively (***Rankin et al., 2009***). We use high-speed cameras, machine-vision analysis, and 300-well plates to quantify habituation across large populations of larvae to identify molecular/genetic mechanisms of long-term habituation (***Figure 1***A,B, ***Randlett et al., 2019***; ***Lamiré et al., 2023***). When stimulated with DFs repeated at 1-minute intervals in blocks of 60 stimuli, larval zebrafish exhibit long-term habituation, reducing not only the probability of executing a response, but also modulating the latency and other kinematic aspects of the response (***Randlett et al., 2019***).

**Figure 1.**
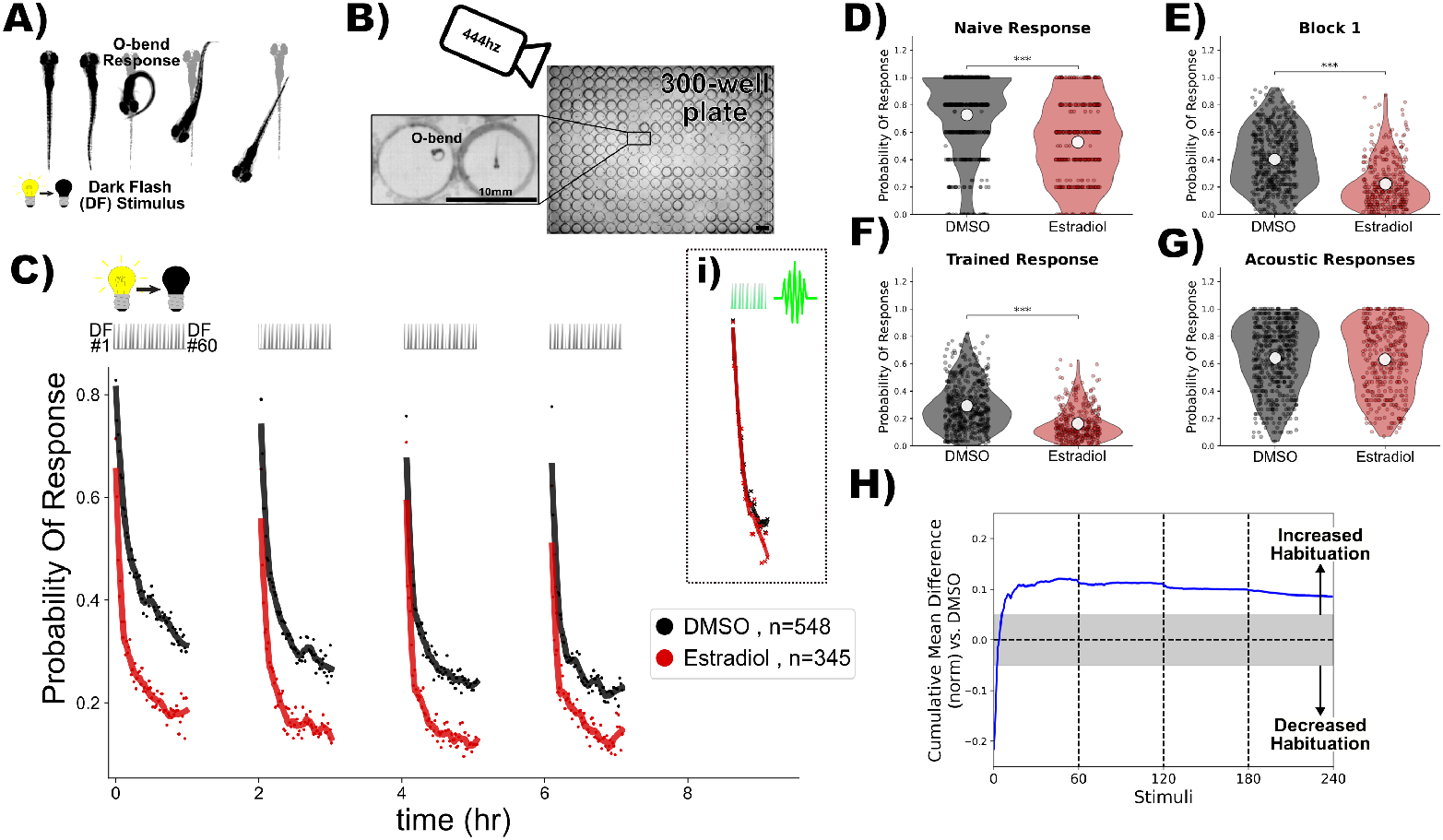
Estradiol increases habituation learning. **A)** In response to a dark flash (DF), larval zebrafish perform a large turning manoeuvre termed an “O-bend” response. **B)** High-throughput setup for recording and quantifying responsiveness using a high-speed camera recording at 444hz observing larvae in 300-well plates. Scale bar = 10mm. **C)** Treatment with estradiol (red) results in more rapid and profound decreases in the probability of response to DF stimuli during habituation training relative to DMSO vehicle controls (black). DF stimuli are delivered at 1-minute intervals, in 4 blocks of 60 stimuli, separated by 1hr of rest (from 0:00-7:00). 1.5 hours later a block of 30 vibration stimuli are delivered at 1-minute intervals (**i**). Each dot is the probability of response to one DF. Lines are smoothed in time with a Savitzky–Golay filter (window = 15 stimuli, order = 2). **D-G)** Distributions responsiveness for different epochs of the experiment. Each dot is the per-fish average of the epoch. Statistical significance was calculated using Mann-Whitney U test, ^***^ = p < 0.001. **D)** the naive response to the first 5 DF stimuli; **E)** the mean response to the remaining DF stimuli in the Block 1 (DFs 6:60); **F)** the trained response to the last 45 DFs in all four training blocks (DFs 16:60,76:120,136:180,196:240); **G)** the 30 vibration stimuli delivered with a tap from a solenoid on the 300-well plate platform. **H)** Cumulative mean difference (CMD) plot quantifying relative habituation performance after estradiol treatment. These plots display the cumulative average differences in the mean response across larvae of the treatment group (estradiol) relative to the control group (DMSO). Difference from 0 reflect a divergence in the change in responsiveness across the 240 DF stimuli in the 4 training blocks, with positive values reflecting increased habituation. The widths of the line is a bootstrapped 99.5% confidence intervals. The gray boxed region reflects the expected non-significant effect size (***Randlett et al., 2019***). Treatment groups are: Estradiol = 10μM estradiol treatment (n = 345 fish); DMSO = 0.1% DMSO vehicle controls (n = 548 fish)

Our previous small-molecule screening experiments identified multiple synthetic Estrogen Receptor agonists as positive modulators of DF habituation learning, including ethinyl estradiol, estradiol valerate, and hexestrol (***Lamiré et al., 2023***). The major effect we observed was a stronger decrease in the probability of executing a O-bend response during the training/learning blocks. We have confirmed and extended these results using estradiol, which is the major natural estrogen in vertebrates.

An acute dose of 10μM estradiol potently increases habituation learning, which is observable when the response probability of the population of estradiol-treated larvae is compared with DMSO-treated vehicle controls (***Figure 1***C-H). Consistent with our previous experiments (***Lamiré et al., 2023***), there is a reduction in the naive responsiveness of the estradiol-treated larvae to the first DF stimuli (***Figure 1***D), but the major effect is observed during the training phase (***Figure 1***C,E,F), as is revealed by the consistent positive deviation in the cumulative mean difference (CMD) plots that are normalized to the naive response level in order to quantify response suppression indicative of habituation performance (***Figure 1***H, ***Randlett et al., 2019***). Importantly, the responsiveness of the larvae to vibration stimuli delivered after the DF stimulation (***Figure 1***Ci), which elicit c-bend escape responses (***Kimmel et al., 1974***), is indistinguishable from controls (***Figure 1***G). From this we conclude that estradiol does not affect global arousal levels but rather has specific effects on habituation learning.

### Gper1 is dispensable for estradiol-promoted habituation learning

The effects of estradiol that we have observed occur very rapidly – larvae are only pretreated with estradiol for ≈25min-1hr before the first DF. This is the time necessary to set the apparatus and begin the experiment. Since the nuclear hormone receptors are thought to primarily exert their effects via transcriptional alterations, this necessitates a delay in their signaling. For this reason, we first hypothesized that membrane-initiated signaling through Gper1 was the most likely mechanism.

To test this we used the *gper1*^*uab102*^ mutant (***Romano et al., 2017***), and generated larvae from *gper1*^*uab102*^ heterozygous or homo/heterozygous crosses to generate clutches of larvae of mixed genotypes. Larvae were treated with estradiol during habituation, and were subsequently genotyped. We reasoned that if *gper1* is required for the effect of estradiol on habituation, mutants would be insensitive to estradiol and habituate significantly less than sibling controls. Contrary to this hypothesis, we found that *gper1* mutants showed no deficits in habituation (***Figure 2***). Remarkably, rather than observing the anticipated inhibition of habituation, *gper1* mutants appeared to habituate slightly more than controls, with the responsiveness level slightly but consistently below the sibling controls across stimuli (***Figure 2***A). This is further supported by a weak but statistically significant decrease in the responsiveness of the larvae during the training period (***Figure 2***D), and a deviation towards positive values in the CMD plot (***Figure 2***F). From these experiments we conclude that Gper1 agonism does not promote habituation learning, but rather may act to suppress it.

**Figure 2.**
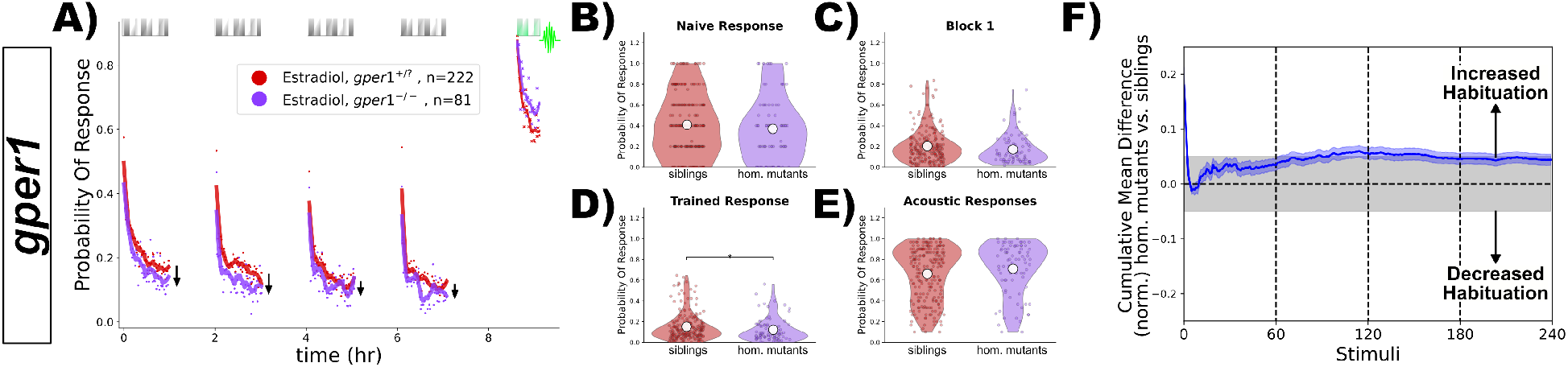
*gper1* mutants do not show habituation deficits after treatment with estradiol. **A)** Homozygous *gper1*^(−*/*−)^ mutants (n = 81 fish, purple) do not show impaired habituation relative to sibling controls (*gper1*^*(+/*−)^ and *gper1*^*(+/+)*^, n = 222 fish, red). Rather, there is a slight suppression of responsiveness in the mutant group (arrows), indicating weakly increased habituation. Each dot is the probability of response to one stimulus. Lines are smoothed in time with a Savitzky–Golay filter (window = 15 stimuli, order = 2). **B-E)** No significant differences are observed in the responsiveness distributions for the naive response to the first 5 DF stimuli (**B**), during the first training block (**C**), or the vibration response (**E**), while a subtle but statistically significant decrease in responsiveness is observed in the trained response (**D**). Statistical significance was calculated using Mann-Whitney U test, ^*^ = p < 0.05. **F)** Cumulative mean difference (CMD) plot quantifying habituation performance of mutants relative to sibling controls, consistent with slightly increased habituation rate in mutant larvae.

### Nuclear ERs are dispensable for estradiol-promoted habituation learning

Since we found that *gper1* was unnecessary for the habituating-promoting effects of estradiol, we next focused on the three nuclear receptors in the zebrafish genome: *esr1, esr2a* and *esr2a*. Using the same strategy as for *gper1*, we analyzed previously established mutants (*esr2a*^*uab134*^, *esr2a*^*uab134*^, and *esr2b*^*uab127*^), looking for a mutant with insensitivity to estradiol. However, we failed to identify any deficits in habituation (***Figure 3***). To our surprise, we again found that both *esr1* and *esr2a* mutants showed subtle increases in habituation (***Figure 3***Avi,Bvi), similar in magnitude to what we had seen for *gper1* mutants (***Figure 2***F). From these data we conclude that none of the nuclear ERs are required for the effects of estradiol on promoting habituation. As for Gper1, ER1 and ER2a show a weak inhibitory effect on habituation, indicating that they also act to suppress habituation learning.

**Figure 3.**
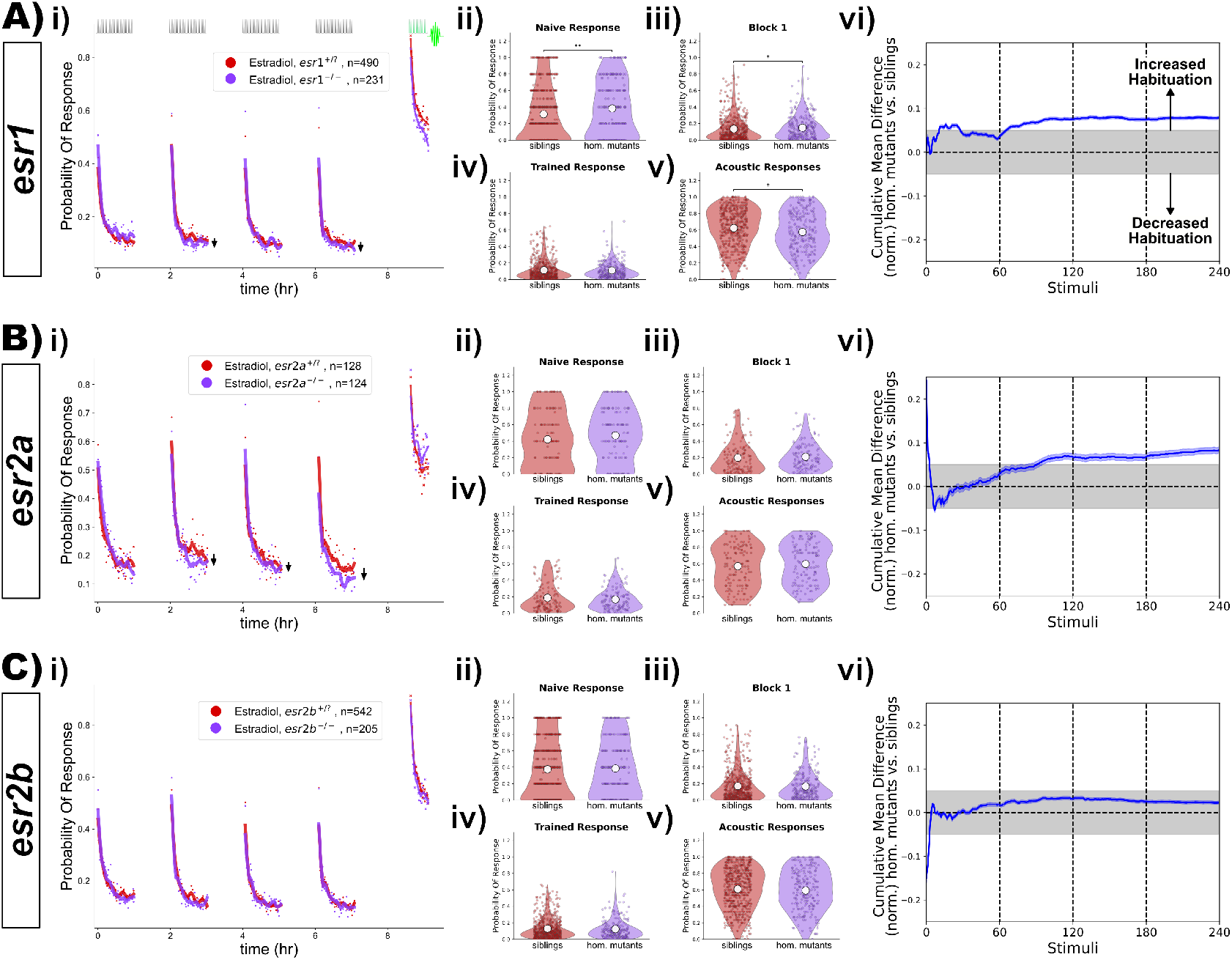
*esr1, esr2a* and *esr2a* mutants do not show habituation deficits after treatment with estradiol. **A)** Homozygous *esr1*^(−*/*−)^ mutants (n = 231 fish) do not show impaired habituation relative to sibling controls (*esr1*^*(+/*−)^ and *esr1*^*(+/+)*^, n = 490 fish). **B)** Homozygous *esr2a*^(−*/*−)^ mutants (n = 214 fish) do not show impaired habituation relative to sibling controls (*esr2a*^*(+/*−)^ and *esr2a*^*(+/+)*^, n = 128 fish). **C)** Homozygous *esr2b*^(−*/*−)^ mutants (n = 205 fish) do not show impaired habituation relative to sibling controls (*esr2b*^*(+/*−)^ and *esr2b*^*(+/+)*^, n = 542 fish). **i)-vi)** For each lettered section: **i)** Responsiveness to stimuli comparing homozygous mutants to sibling controls (heterozygous or wild-type). Each dot is the probability of response to one stimulus. Lines are smoothed in time with a Savitzky–Golay filter (window = 15 stimuli, order = 2). Suppression of responsiveness is indicated by arrows, potentially reflecting increased habituation. **ii)-v)** Distributions responsiveness for different epochs of the experiment. Each dot is the per-fish average of the epoch. Statistical significance was calculated using Mann-Whitney U test, ^*^ = p < 0.05, ^**^ = p < 0.01. **ii)** the naive response to the first 5 DF stimuli; **iii)** the mean response to the remaining DF stimuli in the Block 1 (DFs 6:60); **iv)** the trained response to the last 45 DFs in all four training blocks (DFs 16:60,76:120,136:180,196:240); **v)** the 30 vibration stimuli delivered with a tap from a solenoid on the 300-well plate platform. **vi)** Cumulative mean difference (CMD) plot quantifying habituation performance of mutants relative to sibling controls

### Nuclear ERs are simultaneously dispensable for estradiol-promoted habituation learning

While our experiments demonstrated that *esr1, esr2a* and *esr2a* mutants remain sensitive to estradiol, it is plausible that they could act in a redundant fashion to mediate the effects of estradiol on habituation, perhaps via co-expression in a critical cell type, or via genetic compensation (***El-Brolosy et al., 2019***). To test for this possibility, we generated combinations of mutants by crossing individual lines together (***Figure 4***). A likely scenario could be that the two ERβ paralogs, ER2a and ER2b, act redundantly. However, we found that double mutants for *esr2a;esr2b* did not show habituation deficits (***Figure 4***A). Neither did we observe suppression of habituation in double mutants for *esr1;esr2a* (***Figure 4***B), or *esr1;esr2b* (***Figure 4***C). Finally, we tested triple mutants for all three nuclear ERs (*esr1;esr2a;esr2b*), but again failed to identify suppressions in habituation (***Figure 4***D).

**Figure 4.**
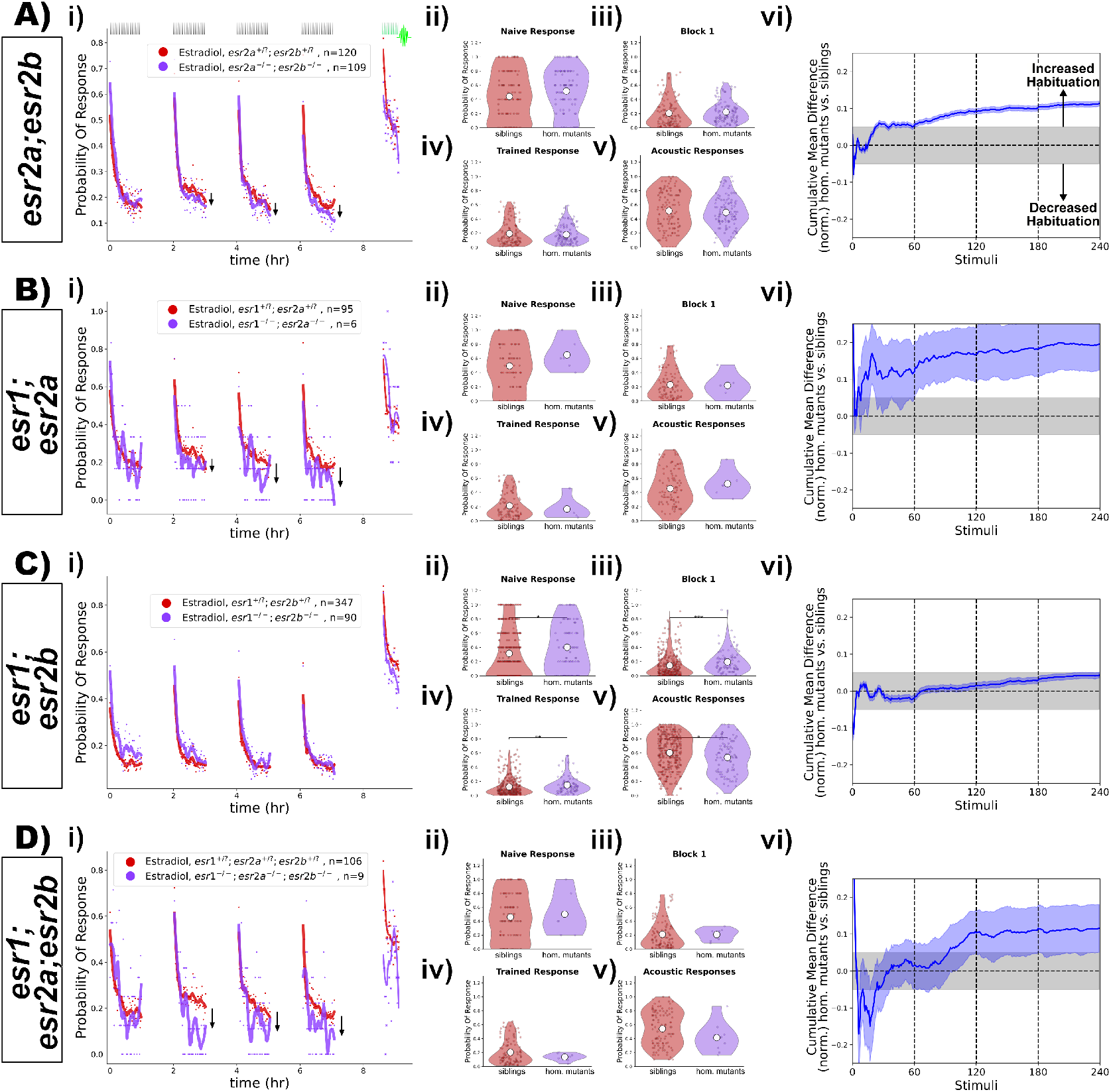
Double and triple mutant combinations of *esr1, esr2a* and *esr2a* do not show habituation deficits after treatment with estradiol. **A)** Homozygous *esr2a*^(−*/*−)^;*esr2b*^(−*/*−)^ double mutants (n = 109 fish) do not show impaired habituation relative to double heterozygous or homozygous sibling controls (^*(+/*?)^;^*(+/*?)^, n = 120 fish). **B)** Homozygous *esr1*^(−*/*−)^;*esr2a*^(−*/*−)^ double mutants (n = 6 fish) do not show impaired habituation relative to double heterozygous or homozygous sibling controls (^*(+/*?)^;^*(+/*?)^, n = 95 fish). **C)** Homozygous *esr1*^(−*/*−)^;*esr2b*^(−*/*−)^ double mutants (n = 90 fish) do not show impaired habituation relative to double heterozygous or homozygous sibling controls (^*(+/*?)^;^*(+/*?)^, n = 347 fish). **D)** Homozygous *esr1*^(−*/*−)^;*esr2a*^(−*/*−)^;*esr2b*^(−*/*−)^ triple mutants (n = 9 fish) do not show impaired habituation relative to triple heterozygous or homozygous sibling controls (^*(+/*?)^;^*(+/*?)^;^*(+/*?)^, n = 106 fish). **i)-vi)** For each lettered section: **i)** Responsiveness to stimuli comparing homozygous mutants to sibling controls (heterozygous or wild-type). Each dot is the probability of response to one stimulus. Lines are smoothed in time with a Savitzky–Golay filter (window = 15 stimuli, order = 2). **ii)-v)** Distributions responsiveness for different epochs of the experiment. Each dot is the per-fish average of the epoch. Statistical significance was calculated using Mann-Whitney U test, ^*^ = p < 0.05, ^**^ = p < 0.01. **ii)** the naive response to the first 5 DF stimuli; **iii)** the mean response to the remaining DF stimuli in the Block 1 (DFs 6:60); **iv)** the trained response to the last 45 DFs in all four training blocks (DFs 16:60,76:120,136:180,196:240); **v)** the 30 vibration stimuli delivered with a tap from a solenoid on the 300-well plate platform. **vi)** Cumulative mean difference (CMD) plot quantifying habituation performance of mutants relative to sibling controls.

We note that we did see a statistically “significant” increase in DF responsiveness in the *esr1;esr2b* double mutants (***Figure 4***Cii-iv). This is attributable to a small increase in naive responsiveness, and did not result from habituation deficits according to our normalized CMD analysis (***Figure 4***Cvi). Considering that we did not observe any increased responsiveness in the triple mutants (***Figure 4***D), we conclude that the observed increased responsiveness in the double mutants is likely a result of biological noise, that only rose to statistical “significance” due to the very large number of larvae tested in our high-throughput experiments.

Consistent with the paradoxical effect of increased habituation in *esr1* and *esr2a* single mutants (***Figure 3***A,B), we again observed that double and triple mutants containing these genes also showed a slight increase in habituation (with the exception of the esr1;esr2b double mutants). This adds further support to the model in which ER1 and ER2b act to suppress learning in this context, rather than promote it.

### Gper1 and nuclear ERs are simultaneously dispensable for estradiol-promoted habituation learning

While it is unclear how the GPCR Gper1 might act redundantly with the nuclear ERs, we nevertheless decided to test this possibility by combining the *gper1* and the *esr1, esr2a*,and *esr2b* mutations (***Figure 5***). As with the previous iterations of this experiment, we did not find combinations of mutants with suppressed habituation (***Figure 5***). Consistent with our model of (*gper1, esr1* and *esr2a*) having inhibitory affects on learning, we again found that most of these combinations of mutants showed evidence of increased habituation (***Figure 5***i,vi). While the responsiveness distributions did not show significant differences (***Figure 5***ii-iv), the normalized CMD plots consistently showed positive deviations, which reflect increased habituation (***Figure 5***vi).

**Figure 5.**
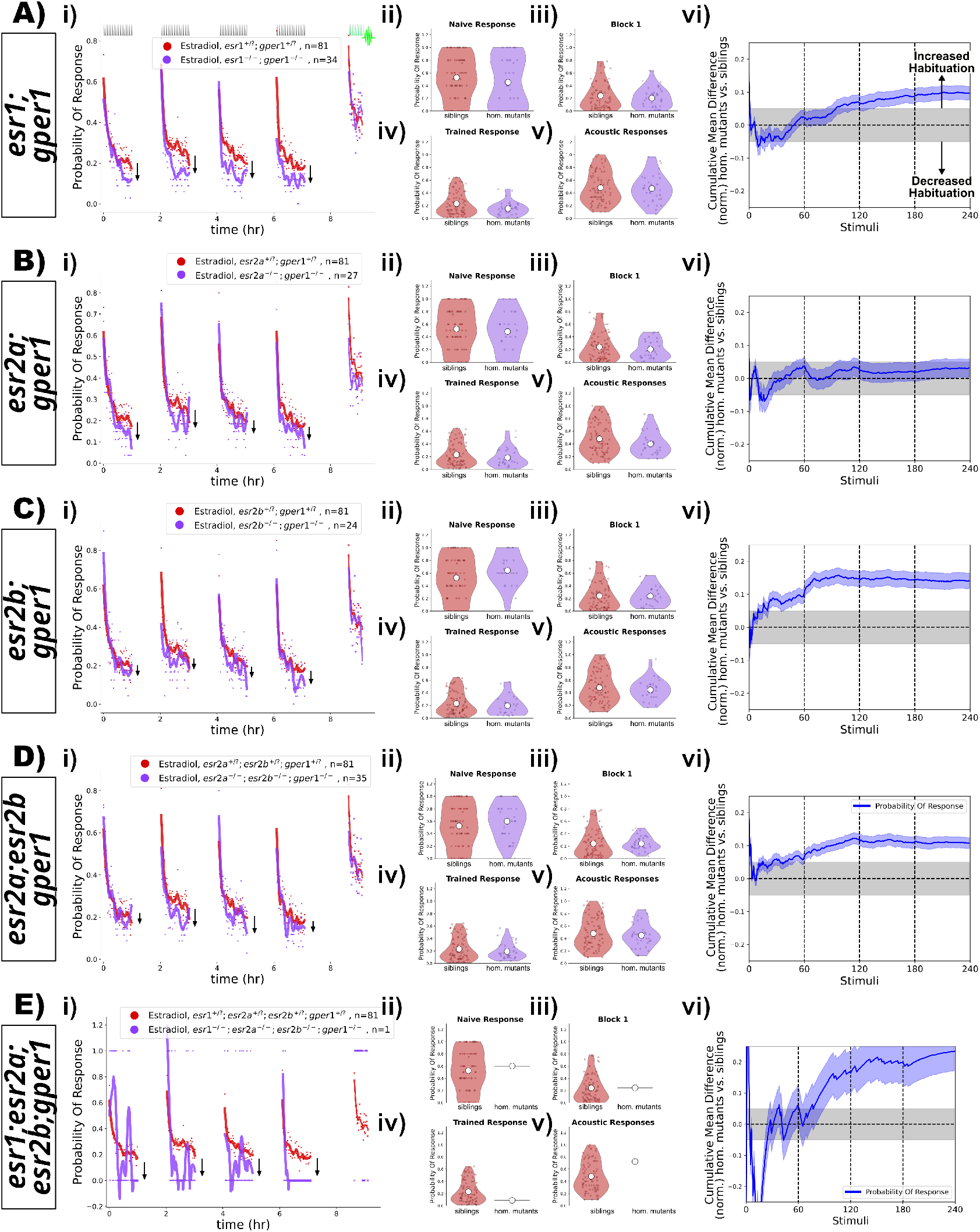
Double, triple and quadruple mutant combinations of *esr1, esr2a, esr2a*, and *gper1* do not show habituation deficits after treatment with estradiol. **A)** Homozygous *esr1*^(−*/*−)^;*gper1*^(−*/*−)^ double mutants (n = 34 fish, purple) do not show impaired habituation relative to double heterozygous or homozygous sibling controls (n = 81 fish, red). **B)** Homozygous *esr2a*^(−*/*−)^;*gper1*^(−*/*−)^ double mutants (n = 27 fish, purple) do not show impaired habituation relative to double heterozygous or homozygous sibling controls (n = 81 fish, red). **C)** Homozygous *esr2b*^(−*/*−)^;*gper1*^(−*/*−)^ double mutants (n = 24 fish, purple) do not show impaired habituation relative to double heterozygous or homozygous sibling controls (n = 81 fish, red). **D)** Homozygous *esr2a*^(−*/*−)^;*esr2b*^(−*/*−)^;*gper1*^(−*/*−)^ triple mutants (n = 35 fish, purple) do not show impaired habituation relative to heterozygous or homozygous sibling controls (n = 81 fish, red). **E)** A single homozygous *esr1*^(−*/*−)^;*esr2a*^(−*/*−)^;*esr2b*^(−*/*−)^;*gper1*^(−*/*−)^ quadruple mutant (n = 1 fish, purple) does not appear to show impaired habituation relative to heterozygous or homozygous sibling controls (n = 81 fish, red). **i)-vi)** For each lettered section: **i)** Responsiveness to stimuli comparing homozygous mutants to sibling controls (heterozygous or wild-type). Each dot is the probability of response to one stimulus. Lines are smoothed in time with a Savitzky–Golay filter (window = 15 stimuli, order = 2). **ii)-v)** Distributions responsiveness for different epochs of the experiment. Each dot is the per-fish average of the epoch. Statistical significance was calculated using Mann-Whitney U test, ^*^ = p < 0.05, ^**^ = p < 0.01. **ii)** the naive response to the first 5 DF stimuli; **iii)** the mean response to the remaining DF stimuli in the Block 1 (DFs 6:60); **iv)** the trained response to the last 45 DFs in all four training blocks (DFs 16:60,76:120,136:180,196:240); **v)** the 30 vibration stimuli delivered with a tap from a solenoid on the 300-well plate platform. **vi)** Cumulative mean difference (CMD) plot quantifying habituation performance of mutants relative to sibling controls. As mutant larvae were all derived from the same experiments, the sibling control data (red) is the same in **A-E)**.

Despite the fact that we analyzed the behaviour of 1152 larvae and successfully genotyping all 4 ER genes in 373 individuals (after >4600 genotyping PCRs), we were only able to identify a single quadruple mutant larva lacking all known ERs (***Figure 5***E). This is likely simply due to the limitations of combinatorial Mendelian inheritance, and the fact that *esr1* and *esr2a* are linked on chromosome 20. While it is dubious to conclude much from an n = 1 experiment, we find it remarkable that this larva exhibits the strongest increased habituation of all of our experiments, with suppression of responses (***Figure 5***Ei), and strong positive deviation in the CMD plot (***Figure 5***Evi).

## Discussion

### The promotion of habituation learning by estradiol is mediated by an unidentified target

Our experiments indicate that Gper1, ER1, ER2a and ER2b do not mediate the positive effects of estradiol on habituation learning. As this is fundamentally a negative result, it is difficult to conclusively demonstrate this beyond any doubt. One major caveat relates to the actual functional nature of the mutant alleles that we have used. These are all Cas9-generated small deletions resulting in frameshift mutations that lead to early stop codons, and are thus predicted null/knockout lines. Despite this genetic confidence, it is always possible that residual activity could still remain, perhaps via alternate splicing or alternate start codons. This could be further complicated by genetic/transcriptional compensation, where frameshift alleles can lead to the upregulation of paralogs in some circumstances (***El-Brolosy et al., 2019***). For this reason, we were originally unsure if we would attempt to publish these results at all. However, we recognize that this bias against publishing negative results is bad for science. Negative results from well-designed and executed experiments are of value for the community and making this knowledge public is our duty as responsible scientists (***Mlinarić et al., 2017***). These high-throughput neurogenetic experiments are also a massive amount of work – raising, testing and genotyping thousands of zebrafish larvae. This was undertaken primarily by technicians and Master’s students, all of whom deserve recognition, irrespective of the perceived “impact” of the outcome of their experiments.

While the possibility of “residual activity” in our mutants is a clear limitation of our approach, we argue that this alternative interpretation is very unlikely. The *esr1*^*uab118*^ and *esr2b*^*uab127*^ alleles both exhibited a lack of estradiol responsiveness in other tissues (***Romano et al., 2017***), and *esr2b*^*uab127*^ mutants are female sterile/subfertile (D. Gorelick, personal communication), indicating a non-functional receptor. Similarly, *gper1*^*uab102*^ mutants show a lack of estradiol responsiveness in heart rate modulation (***Romano et al., 2017***). Interestingly this was only observed in maternal-zygotic mutants. While it seems unlikely that sufficient maternal mRNA/protein for Gper1 could persist in 5dpf larvae, we can formally rule this out with our current datasets. The *esr2a*^*uab134*^ mutants have no previously published phenotype, and so we do not have an independent positive control for the nature of this allele. However, the best evidence we have against the “residual activity” hypothesis is that we actually found phenotypes in our assays for *esr1*^*uab118*^, *esr2a*^*uab134*^, and *gper1*^*uab102*^ mutants. These phenotypes are just of the unexpected sign, where mutants show increased habituation (discussed below).

### Candidate estradiol targets that could promote habituation learning

We have concluded that the lack of habituation deficits in our mutants is due to the presence of an alternative receptor or pathway that mediates the learning-promoting effects of estradiol. In support of this model, a very recent study of estradiol-induced thrombosis came to the same conclusion, namely that an unidentified target mediates this process in larval zebrafish (***Yu et al., 2024***). Importantly, these thrombosis experiments were performed on different knockout alleles which are full genetic deletions, in which “residual activity” is not plausible. Together with our observations, these data suggest that a novel estradiol target exists which has multiple critical functions (at least in zebrafish).

What might this unidentified target be? Various leads exist in the literature. One hypothesis posits the existence of an unidentified “Gq-mER” (Gq-coupled membrane estrogen receptor) (***Qiu et al., 2006***; ***Vail and Roepke, 2019***), and therefore estradiol may signal via additional GPCRs beyond Gper1. Another possibility is an interaction between estradiol and other membrane receptors, for example: the Voltage-Gated Sodium Channel Nav1.2 (***Sula et al., 2021***; ***Treviño and Gorelick, 2021***), transient receptor potential (TRP) channels (***Payrits et al., 2017***; ***Ramírez-Barrantes et al., 2020***), or various other ion channels (***Kow and Pfaff, 2016***). The robust nature of our “non-canonical” but clearly estradiol-dependent phenotype, combined with the high-throughput nature of our behavioural assays, could be an ideal assay for future screening efforts to attempt to identify novel estradiol target(s).

### Multiple Estrogen Receptors act to suppress habituation learning

While we were surprised to find that the classical ERs do not promote habituation, we were shocked to find evidence of the opposite! We found that *esr1*^*uab118*^, *esr2a*^*uab134*^, and *gper1*^*uab102*^ mutants habituate more than their sibling controls (***Figure 1, Figure 2***), consistent with a role for these ERs in inhibiting habituation. While effect sizes of these magnitudes border on those that are easily dismissible as “noise”, they were not only observed in the single mutants experiments, but also in the double and triple mutant combinations of these alleles (***Figure 3***-***Figure 5***), providing good evidence that they are biologically meaningful effects. In fact, these multi-mutants generally exhibited larger effect sizes, consistent with an additive interaction.

Untangling the mechanisms of ER1-, ER2a- and Gper1-dependent suppression of habituation will require considerable further work. The additive interaction we observed genetically indicates that the ERs act cooperatively to suppress habituation learning. All three receptors are expressed in the larval zebrafish brain (***Thisse and Thisse, 2008***; ***Romano et al., 2017***), but whether they are acting in the same or different cell types awaits characterization. The study of this inhibitory pathway may be challenging since it opposes the major learning-promoting effect of estradiol, and therefore may be more straightforward to study after the identification and deletion of the estradiol target that promotes habituation. One attractive hypothesis relates to the observation that estradiol exposure increases aromatase expression in the zebrafish brain (***Menuet et al., 2005***; ***Hao et al., 2013***). Since aromatase catalyzes the conversion of androgens into estrogens, ER mutants may have reduced levels of endogenous endogeous estrogens in the brain, which could lead to inhibited habituation.

## Conclusion

What began as a straightforward study to identify the receptor(s) that mediate the habituation-promoting effects of estradiol has instead led us to a surprising and paradoxical result; canonical ERs do regulate habituation, but are suppressive and act in opposition to the habituation-promoting effects of estradiol. This fits with the general theme of our studies of this habituation paradigm – we find increasing complexity and contradiction within this “simple” learning process the deeper we look. This began with our detailed observations of behaviour, leading us to conclude that habituation results from a distributed plasticity process that adapts different aspects of behavior independently (***Randlett et al., 2019***). We believe that this property underlies our subsequent discoveries of pharmacological and genetic manipulations that can result in either specific changes in specific aspects of habituation (but not others), or even opposing effects, where a single manipulation can simultaneously increase and decrease habituation, depending on which component of behavior is measured (***Randlett et al., 2019***; ***Lamiré et al., 2023***). This complexity appears to be a fundamental property of habituation (***McDiarmid et al., 2019***), and that the study of habituation will likely continue to surprise us, hopefully leading to unexpected insights into the nature of plasticity underlying learning and memory.

## Materials and Methods

### Animals

All experiments were performed on larval zebrafish at 5 days post fertilization (dpf), raised at a density of ≈1 larvae/mL of E3 media supplemented with 0.02% HEPES pH 7.2. Larvae were raised in a 14:10h light/dark cycle at 28-29°C. Adult zebrafish were housed, cared for, and bred at the following zebrafish facilities: Plateau de Recherche Expérimentale en Criblage In Vivo (PRECI, SFR Biosciences, Lyon) and the Animalerie Zebrafish Rockefeller (AZR, SFR Santé Lyon Est, Lyon). Adult zebrafish used to generate larvae were housed in accordance with regulations of the PRECI and AZR facilities, which are regulated by an internal animal wellbeing committee, and were approved by the animal welfare committee (comité d’éthique en expérimentation animale de la Région Rhône-Alpes: CECCAPP, Agreement # C693870602). Behaviour experiments were performed at the 5dpf stage, and are thus not subject to ethical review, but these procedures do not harm the larvae.

Mutant lines were obtained from D. Gorelick’s lab, and were of the following alleles:

*esr1*^*uab118*^ is a 4bp deletion (ZDB-ALT-180420-2), yielding a predicted null frameshift/stop mutation, confirmed by a lack of estradiol responsiveness in the heart as assayed by *Tg(5xERE:GFP)*^*c262*^ expression (***Romano et al., 2017***).

*esr2a*^*uab134*^ is a 2bp deletion (ZDB-ALT-180420-3), yielding a predicted null frameshift/stop mutation (***Romano et al., 2017***)

*esr2b*^*uab127*^ is a 4bp deletion (ZDB-ALT-180420-4), yielding a predicted null frameshift/stop mutation, confirmed by a lack of estradiol responsiveness in the liver as assayed by *Tg(5xERE:GFP)*^*c262*^ expression (***Romano et al., 2017***).

*gper1*^*uab102*^ is a 133bp deletion (ZDB-ALT-180420-1), yielding a predicted null frameshift/stop mutation, confirmed by a lack of estradiol responsiveness in heart beating rate in maternal-zygotic mutants (***Romano et al., 2017***).

### Genotyping

*esr1*^*uab118*^ was genotyped by PCR using the forward/reverse primer pair: GCTGGTCACCTTGAATGCTT/TGAGATGTGAGAGATGACTAGGA with a T_M_ of 58°C yielding a 381 bp PCR product that was digested with the restriction enzyme ApeKI. The mutant product is not digested, and the wild type has two bands at 177 and 204 bp.

*esr2a*^*uab134*^ was genotypes by PCR using the forward/reverse primer pair:

CTTCAGCTGCAGGAAGTGGA/AAAGTCGGGCTTAGCGACTG with a T_M_ of 58°C yielding a 236 bp PCR product that was digested with the restriction enzyme MboI. The mutant product is not digested, and the wild type has two bands at 180 and 56 bp

*esr2b*^*uab127*^ was genotypes by PCR using the forward/reverse primer pair: TGGGCCTGAGATGCAGTAGT/GTGTGTGTCTTGGCCTCCTC with a T_M_ of 60°C yielding a 431 bp PCR product that was digested with the restriction enzyme MbiI. The mutant product is digested into two bands of 150 and 281 bp and the wild type into 3 bands of 78, 150 and 198 bp.

*gper1*^*uab102*^ was genotypes by PCR using the forward/reverse primer pair: ATGGAGGAGCAGACTACCAATGTG/CCATCCAGATGAGGCTGCAA with a T_M_ of 60°C yielding a mutant product of 372bp and a wild type product of 505 bp.

### Pharmacology

β-Estradiol (Sigma E2758, here referred to as “estradiol”) was dissolved in dimethyl sulfoxide (DMSO) and stored at -20°C. Larvae were treated with estradiol immediately before the behavioural assay by pipetting 10-30uL of 10x solution directly into the behavioural wells, always with a final concentration of 0.1% DMSO in E3.

### Habituation behaviour testing

Larval behavior was evaluated in 300-well plates using an updated version of the experimental setup previously described (***Randlett et al., 2019***; ***Lamiré et al., 2023***). Briefly, 300-well plates were custom made using laser-cut acrylic sheets where each well measures 8 mm in diameter and 6 mm in depth, corresponding to an approximate water volume of 300 μL. These plates are suspended under a water bath held at 31°C, acting as a heated lid to minimize condensation and maintain a 29°C water temperature within each well. Behavioral recordings were made using a Mikrotron CXP-4 camera running at 444hz in conjunction with a Silicon Software frame grabber (Marathon ACX-QP, Basler), illuminated by IR LEDs (TSHF5410, digikey.com). Visual stimuli were presented using a rectangular array of 155 WS2813 RGB LEDs (144LED/M, aliexpress.com). For the DF stimulus, the LEDs were briefly switched off (1 s), then linearly returned to the original brightness over a 20 s interval. Vibration/Tap stimuli were administered using a solenoid (ROB-10391, Sparkfun). This behavioral paradigm was designed to be symmetrical: each 1 hr block of stimulation was followed by 1 hr of rest. During these rest periods, the camera was moves using a stepper motor controlled linear actuator (Hanpose HPV4, 500cm), which moved the camera between two plates, allowing us to screen up to 600 fish per experiment across two 300-well plates.

Control of the apparatus (RGB LEDs, solenoid, camera linear actuator) was implemented via a Raspberry Pi Pico microcontroller running CircuitPython (https://circuitpython.org/) (code : code.py) and custom Python software which handled the camera acquisition via the Python wrapper of the Silicon Software Framegrabber SDK, triggered stimuli via the Raspberry Pi Pico, and tracked the head and tail coordinates of the larvae across the 300-wells at a baseline framerate of between 20-30hz (code : Run_BigRig2.py). When a stimulus is delivered (DF or Vibration/Tap), a 1-second “Burst” video is recorded at the full frame rate as a Tiff file, from which the head and tail coordinates are subsequently tracked o?ine (code : TrackBurst_BigRig.py). Larval zebrafish tracking was done via background subtraction and morphological operations implemented using multiple open-source packages, including: OpenCV (***Bradski, 2000***), scikit-image (***Van der Walt et al., 2014***), NumPy (***Harris et al., 2020***), SciPy (***Virtanen et al., 2020***), and Numba (***Lam et al., 2015***).

### Data analysis

Data was analyzed in Python using custom written analysis scripts (code : Analyze_EsrHab.py). Responses to DFs and vibration/taps were identified as movement events that had a cumulative tail bend angle greater than 3 radian (O-bend)and 1 radian (C-bend), respectively. Data was analyzed using multiple open-source packages, including: NumPy (***Harris et al., 2020***), SciPy (***Virtanen et al., 2020***),Numba (***Lam et al., 2015***) and Pandas (***Wes McKinney, 2010***). Data was plotted using Matplotlib (***Hunter, 2007***) and seaborn (***Waskom, 2021***). Statistical “significance” between the distributions was tested using the Mann-Whitney U test implemented in Scipy (***Virtanen et al., 2020***).

The cumulative difference plots to assess changes in habituation performance for the treatments were calculated as previously (***Randlett et al., 2019***), where we first calculated the average response across larvae for each group for each DF. This generated a mean vector for each group. These two vectors were normalized by dividing them by the naive response (mean response to the first 5 DFs), and then the treatment group was subtracted from the control group, yielding a “mean difference” vector between stimulus and controls at each flash. From this mean difference vector we calculated the cumulative mean distribution using Numpy’s ‘nancumsum’ function divided by the number of stimuli experienced, or the index of the vector. To generate statistical confidence in these distributions, we bootstrapped 2000 replicates, and calculated the 99.5% confidence intervals using SciPy’s ‘stats.norm.interval’ function. The assumption of this analysis is that if the two groups are habituating similarly, then the “mean difference” vector will exhibit a noise distribution centered at a mean of 0, and thus the cumulative mean distribution would remain near 0. Treatments that affect habituation will show strong increasing or decreasing cumulative mean distributions, reflecting increased or decreased habituation performance throughout training, respectively. We use an empirically defined threshold of ±0.05 as the statistically meaningful effect size in this analysis, as is reflected in the shaded gray regions in the plots (***Randlett et al., 2019***).

## Data Availability

Software and analysis code is available here: https://github.com/owenrandlett/2025_HabEstrogen. All datasets used in these analyses are available here: HabEstrogen_Datasets.

## Acknowledgments

We thank the Gorelick lab for sharing the ER-mutant zebrafish lines, and for sharing genotyping protocols and expertise, and the Randlett group members for helpful advice regarding the manuscript and work. We are grateful to the staff of the PRECI and AZR zebrafish facilities, including Laure Benard, Robert Renard, Annie Desenfant, Olivier Lohez, for the expert care provided to the zebrafish. We also gratefully acknowledge the communities that develop and maintain the numerous open-source software packages we rely on, most of which could not be cited here.

